# Abnormal weather drives disease outbreaks in wild and agricultural plants

**DOI:** 10.1101/2023.03.17.533130

**Authors:** Devin Kirk, Jeremy M. Cohen, Vianda Nguyen, Marissa L. Childs, Johannah E. Farner, T. Jonathan Davies, S. Luke Flory, Jason R. Rohr, Mary I. O’Connor, Erin A. Mordecai

## Abstract

Predicting effects of climate change on plant disease is critical for protecting ecosystems and food production. Climate change could exacerbate plant disease because parasites may be quicker to acclimate and adapt to novel climatic conditions than their hosts due to their smaller body sizes and faster generation times. Here we show how disease pressure responds to the anomalous weather that will increasingly occur with climate change by compiling a global database (5380 plant populations; 437 unique plant–disease combinations; 2,858,795 individual plant–disease samples) of disease incidence in both agricultural and wild plant systems. Because wild plant populations are assumed to be adapted to local climates, we hypothesized that large deviations from historical conditions would increase disease incidence. By contrast, since agricultural plants have been transported globally, we did not expect the historical climate where they are currently grown to be as predictive of disease incidence. Supporting these hypotheses, we found that disease outbreaks tended to occur during periods of warm temperatures in agricultural and cool-climate wild plant systems, but also occurred in warm-adapted wild (but not agricultural) plant systems experiencing anomalously cool weather. Outbreaks were additionally associated with higher rainfall in wild systems, especially those with historically wet climates. Our results suggest that historical climate affects susceptibility to disease for wild plant–disease systems, while warming drives risks for agricultural plant disease outbreaks regardless of historical climate.

## MAIN TEXT

Infectious disease outbreaks cause massive losses in crop yields^1^, threaten food security^2^ and imperil wild plant populations^3^. Both temperature and moisture exert strong effects on plants and the parasites that infect them^4^ and can be key to accurately predict plant disease across time and space^5^, making these systems highly susceptible to environmental effects across biological scales^6^. Critically, relatively little is known about how plant–disease systems will respond to novel weather caused by climate change^7^, despite the fact that these responses may have implications for global food security, plant conservation, and ecosystem management. To anticipate and mitigate climate change effects, we must understand how past climate and current weather have shaped plant disease around the world.

Animal populations adapted to cold climates experience larger disease outbreaks under unusually warm weather, while animals adapted to warm climates experience more disease under cold weather^8^. A proposed mechanism for these ‘thermal mismatches’ is that, on average, small bodied organisms such as parasites have functionally wider thermal breadths (i.e., the range of temperatures at which an organism has strong performance) than larger bodied organisms^9^, possibly due to quicker acclimation or adaptation. According to this thermal mismatch hypothesis, parasite performance at non-optimal temperatures is expected to be less negatively impacted than host performance^8,10^. One prediction that follows is that plant species adapted to cooler climates may experience greater disease pressure in warm weather, and that warm-adapted plants may conversely experience greater outbreaks in cooler weather. The thermal mismatch hypothesis therefore predicts a negative interaction between historical (climate) and contemporaneous (weather) temperature effects on disease due to the smaller parasites performing relatively better at abnormal temperatures compared to their larger bodied hosts.

Beyond temperature change, other aspects of climate change can affect plant diseases, including changes in CO_2_ concentrations and precipitation^11,12^. Moisture levels have long been known to regulate plant infections^4^, and both drought or extreme high precipitation levels could increase disease under different contexts. Indeed, drought can increase physiological stress and therefore vulnerability to pathogen attack^13^, while extreme precipitation can increase spread of certain plant parasites^14,15^, as well as wash away contact pesticides^16^. It is less clear if plant and parasite adaptation to historical precipitation levels mediate effects of current rainfall and moisture.

Importantly, climate–disease dynamics in agricultural systems may differ from those in wild systems. In agricultural systems, farmers can select for genotypes that are adapted to their new climates^17^, while adaptation in wild systems may be limited by genetic population structure^6^. Additionally, while climate change can cause modest range shifts in wild plants (e.g., < 100 meters/year in beech and maple trees^18^), agricultural plants have been moved around the planet for centuries^19^, often experiencing artificial selection for high yields across environmental gradients. Finally, extreme weather and disease pressure are often mitigated in agricultural systems (e.g., through irrigation, shading, or pesticide application) but not in wild systems. We suggest, therefore, in contrast to wild plants, the historical climate where agricultural plants are currently grown may not be as predictive of their sensitivity to disease under novel weather.

To test the hypothesis that anomalous temperature and precipitation impact disease outbreaks in wild and agricultural plants, we assembled a large population-level database of plant disease incidence in wild and agricultural systems from the literature (n = 5380 plant populations, n = 437 unique host–disease combinations), then paired georeferenced climate and weather data with each observation. We fit a binomial mixed-effects model to the data to test our predictions that contemporaneous weather affects plant disease incidence, that there are negative interactions between climate and weather on plant disease (thermal and/or precipitation mismatches), and that the effects of weather and climate differ between wild and agricultural systems.

### Effects of climate and weather on plant disease incidence

Our database comprised 5380 population-level observations of infectious disease incidence (number of plants infected / number of plants surveyed) in plants, arising from nearly three-million individual plant– disease samples. The data span four broad types of disease-causing agents (Fig. 1a), 29 host plant orders (Fig. 1b), and six continents in the time period 1984–2019 (4622 observations in agricultural systems from 99 studies; 758 observations in wild systems from 17 studies)(Fig. 1c; Table S1).

**Fig 1.**
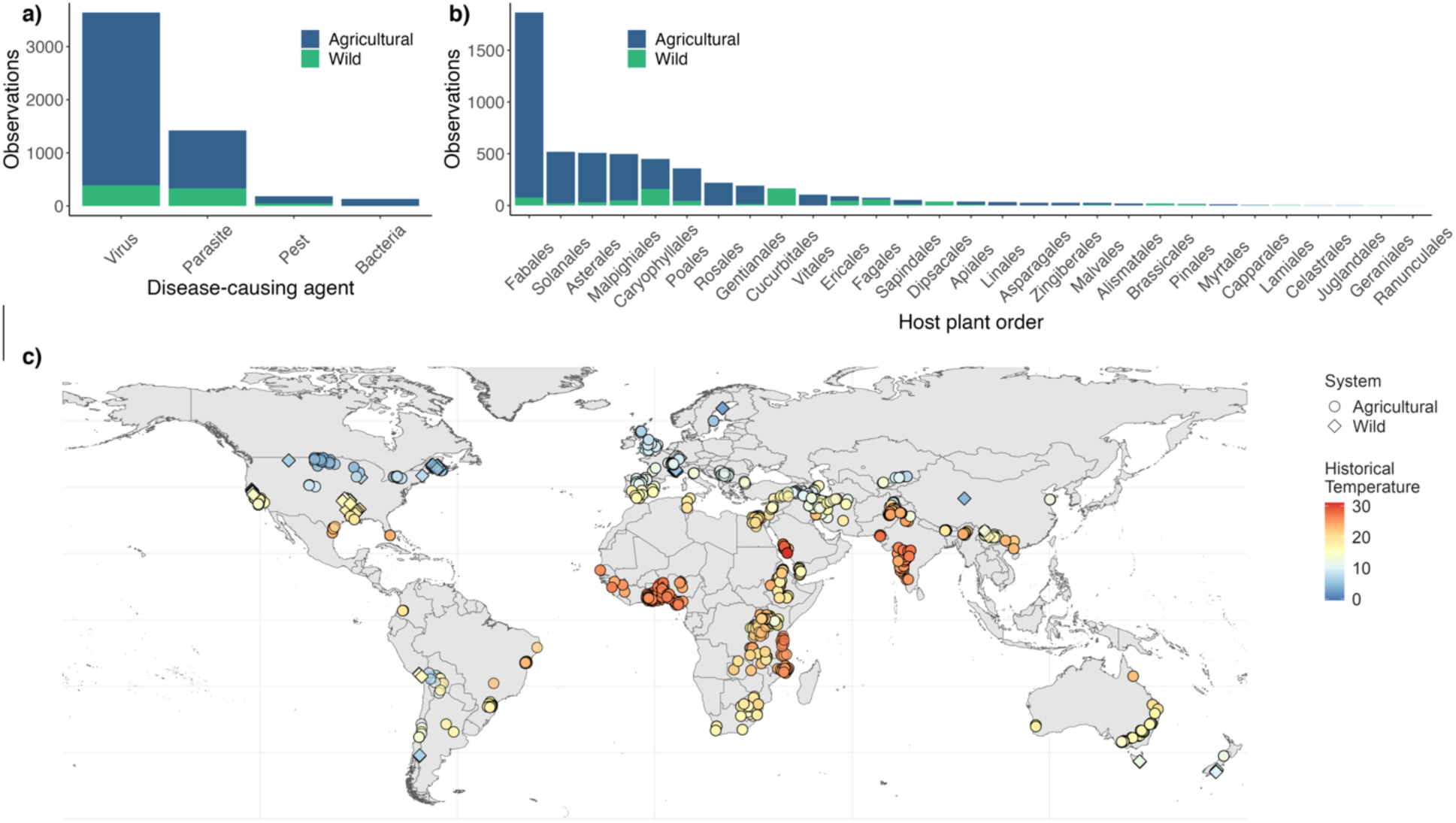
A global database of plant disease and climate spanning geography, climate, parasites, and hosts. a) Number of survey observations by type of disease-causing agent across agricultural (blue) and wild (green) systems. ‘Parasite’ represents eukaryotic parasites. b) Number of survey observations by host plant order across agricultural and wild systems. c) Records of plant disease incidence in agricultural (circles) and wild (diamond) populations (5380 observations). Point color represents the average historical temperature for each location (°C).

We found significant effects of historical climate and contemporaneous weather on plant disease, and these effects differed in strength and direction between agricultural and wild plant systems (Fig. 2). Historical and contemporaneous temperature had, respectively, small and moderate positive effects on disease in agricultural systems but strong negative effects in wild systems (Fig. 2). Importantly, the interaction between historical and contemporaneous temperature was small and positive in agricultural systems but large and negative in wild systems, revealing that thermal mismatches distinctly occur in wild systems (Fig. 2). As expected, precipitation effects on disease were also significantly larger in wild systems than agricultural systems (Fig. 2). Disease risk also varied by the type of disease-causing agent, with pests associated with relatively high disease incidence, viruses and eukaryotic parasites intermediate incidence, and bacteria relatively low incidence (Fig. S3).

**Figure 2.**
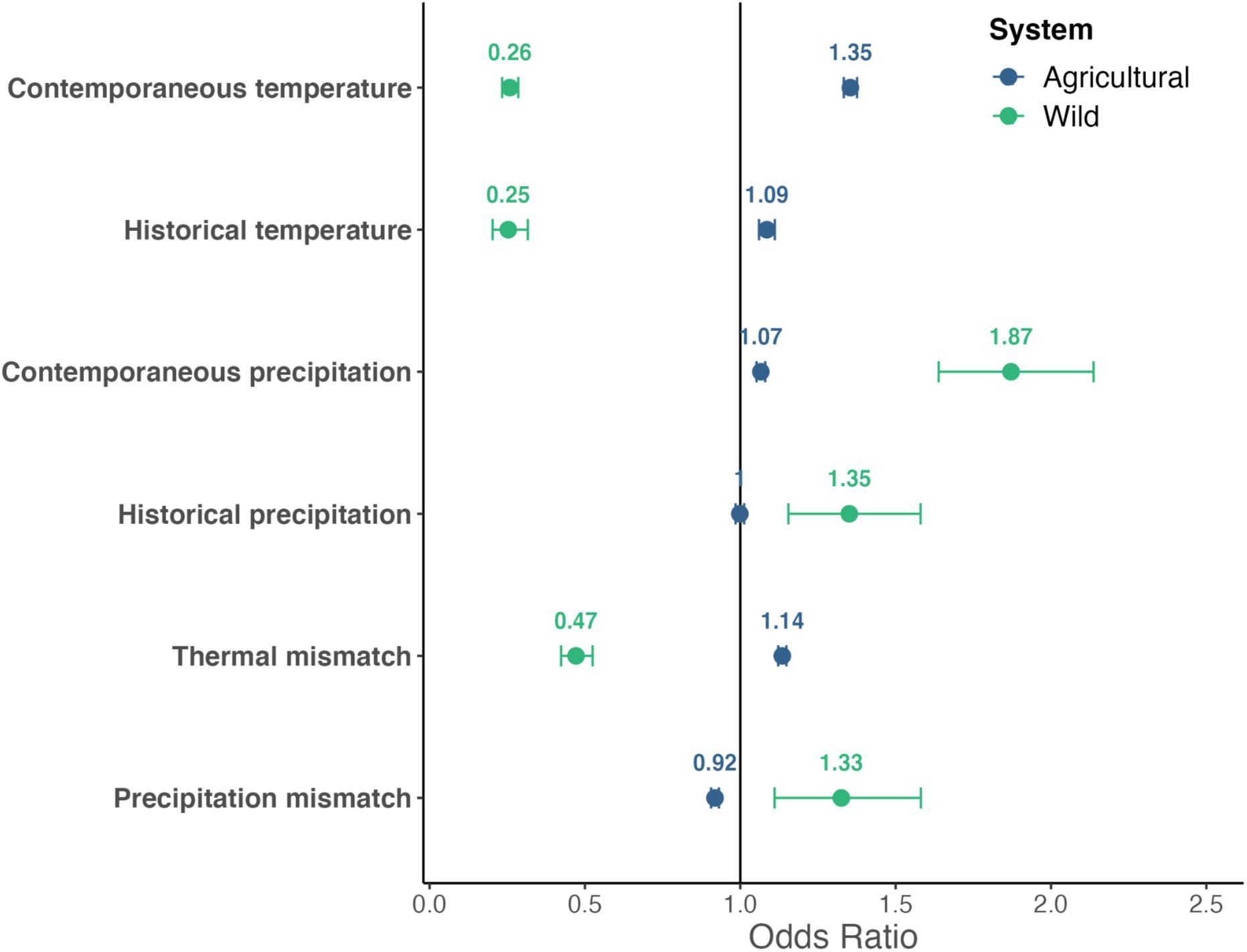
Climate and weather effects on disease differed between agricultural and wild systems, including strong thermal mismatches in wild systems and strong warm weather effects in agricultural systems. Points and error bars represent estimated odds ratios and 95% confidence intervals, respectively, where higher odds ratios represent relatively greater disease risk. Odds ratios for agricultural plants are shown in blue, and for wild plants in green. Contemporaneous and historical temperature and precipitation were standardized before model fitting. Here, thermal/precipitation mismatches represent an interaction between contemporaneous temperature/precipitation by historical temperature/precipitation, respectively, though the mismatch term is typically reserved for *negative* interactions between these variables. Odds ratios associated with the different types of disease-causing agents included in the model are shown in Fig. S3.

### Temperature effects differ between wild and agricultural systems

Effects of temperature on disease were positive in agricultural systems regardless of historical climate (Fig. 3b). In contrast, wild systems exhibited a thermal mismatch in which warming led to greater disease in historically cool wild systems, but historically warm wild systems experienced greater disease at abnormally cool temperatures (Fig. 3a). This interactive effect is consistent with the idea that climate maladaptation is an important driver of disease outbreaks, and parallels findings in animal systems^8,10,20^. Our result lends further support to the thermal mismatch hypothesis, which predicts increased disease at temperatures that deviate from those to which hosts are adapted^10^. Our findings therefore demonstrate that wild plant systems in temperate zones, which are more likely to be cold adapted, are most vulnerable to warming-driven outbreaks, while systems in tropical zones may experience less disease pressure with warming.

**Figure 3.**
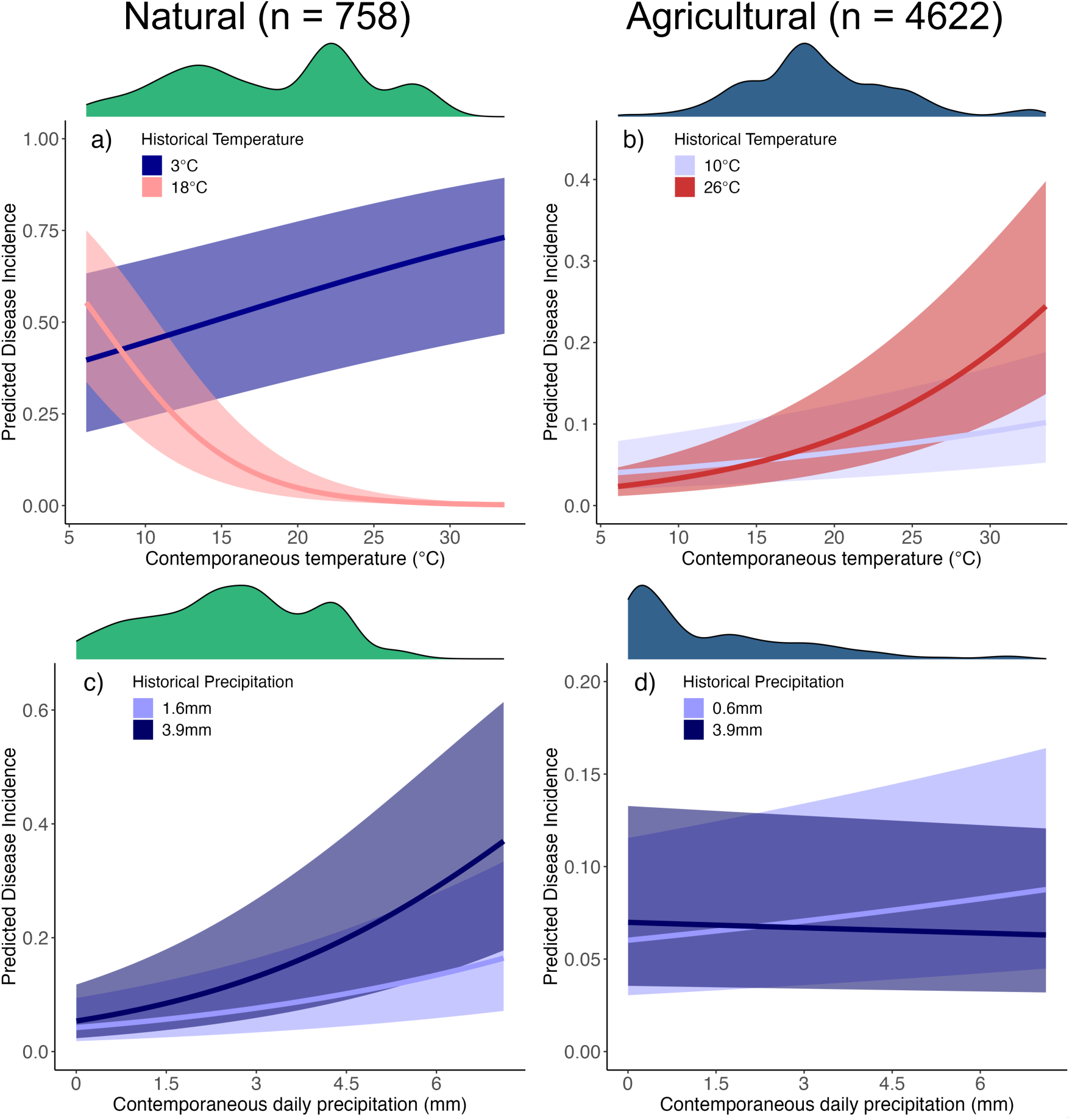
Warm weather promotes disease incidence in agricultural and cool-adapted wild systems, cool weather promotes disease in warm-adapted wild systems (a thermal mismatch). Wet weather promotes disease in wild systems, especially in wetter climates. Lines show the estimated marginal effects of contemporaneous temperature (a, b) and contemporaneous daily precipitation (c, d) on disease incidence in wild (a, c) and agricultural (b, d) systems for the 10% and 90% quantiles of historical temperature and historical precipitation, which differ between agricultural and wild systems. X-axis has been changed from standardized weather metrics to temperature (°C) and daily precipitation (mm) for ease of interpretability. These marginal effects are generated under the condition that the disease-causing agent is a virus, which was the most surveyed. Density plots above each panel represent the distribution of contemporaneous temperature (a, b) and contemporaneous precipitation (c, d).

In contrast to wild plant systems, the evidence suggests that agricultural plant populations from both cold and warm climates experienced greater disease incidence with warming (Figs. 2, 3b). Agricultural plant populations typically have comparatively short evolutionary histories in their present-day locations, and are frequently artificially selected for traits that enhance crop production and other human-preferred characteristics^21^. While crop and forestry species and subspecies varieties are in part chosen based on their ability to grow and produce under a certain climate^17^ and gene flow from wild relatives can increase local crop genetic diversity^19^, we hypothesized that the agricultural systems would still be less adapted to local climatic conditions than wild populations because selection for human-preferred characteristics may not be directly aligned with preferred adaptations to local climate. Thus, it is not surprising that they do not show the thermal mismatch present in wild systems. Instead, warming increased disease incidence on average across all agricultural systems, and these increases were slightly larger for agricultural plants from warm climates compared to those in cooler climates (Fig. 3b). It is possible that farmers in warm climates (e.g., average annual temperature > 20°C) have less capacity to mitigate the consequences of unusual weather (e.g., through watering, shading, or pesticide application) due to differences in agricultural mechanization^22^. While we did not have fine grained data to explore the potential effects of these factors here, it could be beneficial to incorporate socioeconomic data into future work that seeks to make more local-scale predictions of climate effects on agricultural disease.

Our database captures the positive correlation (*r* = 0.56; Fig. S1) between historical and contemporaneous temperature: historically cold places are more likely to have colder weather during disease surveys and vice versa (Fig. S2). Additionally, observations in wild systems typically occurred at cooler temperatures than those in agricultural systems (Fig. 3, Fig. S2). Together, these data features mean that there is weak justification for extrapolating trends observed for cold temperatures to trends expected at warm temperatures in wild systems, or from warm to cold temperatures in agricultural systems. Experimental temperature manipulations that can supplement observational approaches by overcoming these constraints would allow tests of both wild and agricultural plant systems from different historical climates against a range of experimental temperatures. Such experiments may also parse out the relative importance of different mechanisms by which thermal mismatches impact population-level disease incidence, including through effects on plant immunity and resistance, and on parasite growth and transmission.

### Precipitation effects are stronger in wild systems and mediate thermal mismatches

Higher precipitation during survey periods was associated with greater disease incidence, with the strongest effects in wild systems. Precipitation effects differed across system types: in wild systems, contemporaneous rainfall effects were strong, particularly in historically wet locations (Fig. 3c), while in agricultural systems precipitation effects were generally weak (Fig. 3d). Overall, precipitation effects were weaker than temperature effects, a result that has previously been shown experimentally for foliar fungal diseases in a wild alpine meadow system^23^.

The observation in wild systems that contemporaneous rainfall effects were stronger in historically wet locations (Fig. 3c) is surprising from a host perspective because we might expect plants to be best defended against infection under precipitation conditions to which they are adapted (c.f. thermal mismatch hypothesis). However, we lack the data on host and parasite moisture–performance curves, therefore it is unclear if they follow the same body size dependencies observed with thermal performance curves in which parasites have larger breadths than hosts. It is possible that pathogens and parasites from historically wet locations have evolved to transmit under high levels of rainfall (e.g., through splash-mediated dispersal), and that increases in contemporaneous rainfall in these regions benefit the pathogen more than they do the hosts. Alternatively, higher rainfall could lead to more disease because it facilitates the invasion of wet-adapted parasites, or because historically present pathogens that evolved in wetter climates will then perform better as rainfall increases. Collecting moisture–performance curves across host–parasite systems could allow for these different potential mechanisms to be assessed.

Projected effects of climate change on precipitation are more variable than those of temperature, with some areas likely to see large increases in rainfall while other areas experience significant decreases^24^. This variability suggests that plant disease incidence will increase in some areas while decreasing in others, similar to projections for how changing rainfall patterns will alter distribution of human diseases like malaria in West Africa^25^ and cholera across Africa^26^. Projections for how climate change will affect plant diseases will thus need to account for potential differences in how temperature and local precipitation patterns will change relative to each other. For instance, the downy mildew – grapevine system exhibits positive relationships between both temperature and disease and rainfall and disease^14^. Despite projected decreases in rainfall for the mildew – grapevine system’s region of Italy, disease epidemics are expected to increase because increases in temperature-driven disease will more than offset any reduction due to decreased rainfall^14^.

Interactions between past and current temperatures may also interact with precipitation. We tested for these interactions separately in the wild and agricultural systems using modified versions of our main model reported above (see Materials and Methods). This secondary analysis found that thermal mismatches (i.e., negative interactions between contemporaneous and historical temperature) became stronger under wetter conditions in wild systems (Fig. S4). In agricultural systems, we found synergistic effects between the temperature interaction and contemporaneous precipitation (Fig. S5), meaning that agriculture in locations that are projected to increase in both temperature and precipitation may suffer from even greater increases in disease than would be expected based on the sum of their effects. Historical climate may, therefore, be more important for predicting the effects of weather on plant diseases under high rainfall, but less important if rainfall is low. Future models that incorporate the direct and possibly synergistic effects of additional environmental axes (e.g., CO_2_, which can interact with temperature to regulate plant pest populations^27^) are likely to exhibit increased predictive accuracy.

### Conclusions

Understanding how weather anomalies mediate plant disease outbreaks is critical for anticipating and mitigating climate change impacts for agricultural and wild plant systems, which affect food security^2^ and ecosystem integrity^7^. Here, we compile and analyze a novel spatiotemporal plant disease database, revealing that disease outbreaks occur more commonly during periods of warm temperatures in agricultural and cool-climate wild plant systems. However, in warm-climate wild plant systems, anomalously cool weather is associated with larger disease outbreaks. Outbreaks were associated with higher rainfall in wild systems, especially those systems with historically wet climates. Together, these results suggest that the evolutionary history of wild plant – disease systems in their historical climate affects vulnerability to disease. Agricultural systems, which have typically had a shorter evolutionary history in their present-day locations, are vulnerable to pathogen outbreaks under warm conditions regardless of historical climate. Together, these data-driven conclusions offer general rules of thumb that may apply to understudied systems under anomalous weather and climate.

Our work illustrates that warming temperatures are likely to drive outbreaks in many agricultural and historically cool wild plant systems. As climate change-driven precipitation trends are heterogeneous in space^24^, enhanced disease from warming is likely to be compounded by increased rainfall in some areas but mitigated by decreased rain in others. This work allows us to cut through the complexity of plant - disease - climate interactions to understand that effects of climate on disease outbreaks can be predictable based on historical climate.

## Materials and Methods

### Database construction: literature search for plant disease surveys

We conducted a systematic literature review of published plant disease surveys to compile a global, spatiotemporal database of plant disease incidence (number of infected plants / number of plants sampled). We searched the Web of Science via Stanford University library in February, 2021 using combinations of the search terms parasit*, survey*, disease*, pest*, pathogen*, damage*, vir*, plant*, crop*, tree*, forest*, prevalence*, incidence*, percent*, and proportion*, which returned 1800 studies. After compiling our database, we discovered that certain geographic regions—namely, South America, Central America, and the Malay Archipelago—were underrepresented. We thus conducted additional searches in Web of Science using combinations of the search terms above with the names of each country in these regions and screened an additional 582 non-mutually exclusive studies.

We screened abstracts as including potentially relevant data or not, using criteria defined below. We then read each study scored as having potentially relevant data and determined if the study included information on: (1) an approximate location or latitude/longitude coordinates, (2) sample size of the survey, (3) month(s) and year in which the survey took place (up to a maximum of six consecutive months), (4) identity of plant host and disease-causing agent, and (5) either disease incidence (number of infected plants / number of total plants surveyed) or the number of infected samples. We also confirmed that each study used randomly selected samples to calculate incidence and that surveys occurred outdoors (i.e., not in a glasshouse). If information was partly present, we attempted to contact the corresponding authors via email to obtain the missing information. Studies included in our analyses are listed in Table S1.

Extracted data for each disease survey included host and parasite taxonomic information, the month(s) in which the survey occurred, plant sample size and disease incidence, and survey location information. If the study did not include taxonomic information, we searched the colloquial name and used the Integrated Taxonomic Information System to retrieve it. We recorded latitude and longitude if provided by the study, and if this information was not provided we extracted approximate latitude and longitude from Google Maps for the centroid of the named location(s) of the survey. Additionally, we recorded the approximate spatial scale over which a survey occurred by using the ruler tool in Google Maps along with information provided by each study (e.g., “survey occurred throughout X county”) to measure the average distance across the survey range, with a minimum distance cutoff of 1km. Median distance across the dataset was 20km, and the maximum distance was 500km.

### Database construction: climate and weather data

We paired each observation of plant disease incidence with climate reanalysis data extracted using Google Earth Engine^28^. First, we specified circular buffers around the center coordinates of each observation with diameter equal to the approximate spatial distance of the survey. We then extracted temperature and precipitation (weather) data from the ERA5-land monthly averaged dataset^29^ for the location and month(s) that each incidence survey occurred, and calculated the mean temperature and mean daily precipitation over the months of the survey period. We extracted historical average temperature and precipitation data (30-year averages from 1960-1990) for the location of each observation from the WorldClim BIO Variables dataset^30^. These historical data are annual averages, rather than seasonal, because growing season averages could mask the variability between climates that plants need to cope with in colder regions during spring and fall.

### Analyses

We fit a binomial mixed-effects model to the data (5380 populations from 2,858,795 plant–disease samples) using the glmmTMB function from the *glmmTMB* package^31^ in R^32^. This model included fixed effects for historical temperature, contemporaneous temperature, historical precipitation, contemporaneous precipitation, type of disease-causing organism (bacteria, virus, eukaryotic parasite, or animal pest such as an insect or nematode), a factor for agricultural or wild plant hosts, two-way interactions between historical and contemporaneous temperatures (i.e., where a negative interaction constitutes a thermal mismatch), historical and contemporaneous precipitation (i.e., where a negative interaction constitutes precipitation mismatch), each of the four temperature or precipitation predictors interacting with the factor for agricultural or wild plant hosts, as well as a three-way interaction between agricultural or wild plant hosts with historical and contemporaneous temperatures, and a three-way interaction between agricultural or wild plant hosts with historical and contemporaneous precipitation. All weather and climate variables were standardized as z-scores. To control for non-independence among host phylogeny and data collected within each study, the model also included host plant order and study ID as random effects.

We fit similar, adjusted versions of the model above to test additional hypotheses regarding interactive effects of precipitation and temperature. To accommodate the additional interaction terms required, we separated the wild (n=758) and agricultural (n=4622) systems, and tested whether precipitation alters the strength of thermal mismatches by fitting identical models that included a three-way interaction between historical and contemporaneous temperature and contemporaneous precipitation. These models also included each two-way interaction, each of the three main effects, an effect for the type of disease-causing organism, and random effects for host plant order and study ID, as described for the main model above.

## Acknowledgements

We thank members of the Mordecai and O’Connor labs for valuable feedback on the manuscript.

## Author Contributions

All authors conceived of the study. DK and VN conducted the literature review. DK, JC and MC conducted analyses. DK wrote the first draft of the manuscript, and all authors significantly contributed to editing and revising the manuscript.

## Competing Interests

The authors declare no competing interests.

## Notes

### Competing Interest Statement

The authors have declared no competing interest.

