## Supporting Information for "Abnormal weather drives disease outbreaks in wild and agricultural plants"

### Additional details on database construction

*Literature search for plant disease surveys*

As described in the main text, we conducted a systematic literature review of published plant disease surveys to compile a global, spatiotemporal database of plant disease incidence. Our original search in February 2021 using the Web of Science via Stanford University library used combinations of the search terms parasit*, survey*, disease*, pest*, pathogen*, damage*, vir*, plant*, crop*, tree*, forest*, prevalence*, incidence*, percent*, and proportion*, which returned 1800 studies. After compiling our database, we discovered that certain geographic regions—namely, South America, Central America, and the Malay Archipelago—were underrepresented.

We thus conducted additional searches in April 2022 in Web of Science using combinations of the search terms above with the names of each country in these regions, and screened an additional 582 non-mutually exclusive studies. This resulted in the inclusion of seven additional studies from South America, though none of the studies we reviewed for Central America or the Malay Archipelago included all required information or met all criteria. It is likely that surveys with data relevant to the investigations in this study do exist for these and other regions that are underrepresented in our study, either in other databases or published in non-English journals. Future studies that are able to compile survey data across more published languages could potentially fill these important geographical gaps.

Studies included in our analysis are listed in Table S1:

**Table S1. Plant disease survey studies compiled in our novel database.** Studies may appear more than once in Table S1 if they report survey results for more than one type of system, host order, or disease-causing agent. Observations represents the number of population-level surveys from each study that is included in our database.

| Reference | System | Host order | Disease-causing agent | Country of survey(s) | Observations |
|---|---|---|---|---|---|
| (Abbate and Antonovics, 2014) | Natural | Caryophyllales | Eukaryotic parasite | France | 122 |
| (Adediji et al., 2015) | Agricultural | Sapindales | Virus | Nigeria | 11 |
| (Aguayo et al., 2014) | Natural | Fagales | Eukaryotic parasite | France | 13 |
| (Aldhebiani et al., 2015) | Agricultural | Rosales | Virus | Saudi Arabia | 60 |
| (Ale-Agha and Rakhshandehroo, 2014) | Agricultural | Rosales | Virus | Iran | 28 |
| (Abdel Aleem et al., 2018) | Agricultural | Solanales | Virus | Egypt | 56 |
| (Alegbejo et al., 2008) | Agricultural | Malvales | Virus | Nigeria | 6 |
| (Ali and Randles, 1997) | Agricultural | Fabales | Virus | Pakistan | 24 |
| (Ali et al., 2002) | Agricultural | Solanales | Virus | Pakistan | 84 |
| (Alicai et al., 2007) | Agricultural | Malpighiales | Virus | Uganda | 54 |
| (Arain et al., 2012) | Agricultural | Malvales | Eukaryotic parasite | Pakistan | 4 |
| (Ávila et al., 2009) | Agricultural | Solanales | Virus | Brazil | 48 |
| (Banito et al., 2007) | Agricultural | Malpighiales | Bacteria | Togo | 4 |
| (Banito et al., 2007) | Agricultural | Malpighiales | Virus | Togo | 4 |
| (Bao et al., 2007) | Agricultural | Fabales | Virus | China | 48 |
| (Bao et al., 2007) | Natural | Fabales | Virus | China | 2 |
| (Bekele et al., 2005) | Agricultural | Fabales | Virus | Ethiopia | 480 |
| (Biswas et al., 2016) | Agricultural | Sapindales | Virus | India | 27 |
| (Bokonon-Ganta and | Natural | Sapindales | Pest | Benin | 9 |

| | | | | | |
|---|---|---|---|---|---|
| Neuenschwander, 1995) | | | | | |
| (Borza et al., 2019) | Agricultural | Rosales | Eukaryotic parasite | Canada | 4 |
| (Borza et al., 2019) | Agricultural | Solanales | Eukaryotic parasite | Canada | 2 |
| (Brentu et al., 2012) | Agricultural | Sapindales | Eukaryotic parasite | Ghana | 6 |
| (Bressan et al., 2011) | Agricultural | Caryophyllales | Bacteria | France | 24 |
| (Brewer et al., 2012) | Agricultural | Malvales | Bacteria | USA | 10 |
| (Brey et al., 1998) | Natural | Poales | Pest | USA | 35 |
| (Bruez et al., 2013) | Agricultural | Vitales | Eukaryotic parasite | France | 56 |
| (Buzkan et al., 2013) | Agricultural | Solanales | Virus | Turkey, Tunisia | 28 |
| (Carloni et al., 2013) | Agricultural | Poales | Bacteria | Argentina | 4 |
| (Carlsson et al., 1990) | Natural | Caryophyllales | Eukaryotic parasite | Sweden | 15 |
| (Carlsson et al., 1990) | Natural | Dipsacales | Eukaryotic parasite | Sweden | 39 |
| (Carlsson et al., 1990) | Natural | Ericales | Eukaryotic parasite | Sweden | 45 |
| (Castillo and Plata, 2016) | Agricultural | Solanales | Bacteria | Bolivia | 16 |
| (Cheah et al., 2003) | Agricultural | Asparagales | Eukaryotic parasite | Australia | 24 |
| (Chitambo et al., 2019) | Agricultural | Caryophyllales | Pest | Kenya | 12 |
| (Chitambo et al., 2019) | Agricultural | Solanales | Pest | Kenya | 24 |
| (Chittem et al., 2015) | Agricultural | Fabales | Eukaryotic parasite | USA | 38 |
| (Coelho et al., 2021) | Agricultural | Sapindales | Pest | Brazil | 1 |
| (Daugherty et al., 2018) | Agricultural | Vitales | Bacteria | USA | 6 |
| (de Oliveira et al., 2020) | Agricultural | Malpighiales | Eukaryotic parasite | Brazil | 17 |
| (Perez De San Roman et al., 1996) | Agricultural | Caryophyllales | Virus | Spain | 107 |

| | | | | | |
|---|---|---|---|---|---|
| (Elbeshehy et al., 2017) | Agricultural | Gentianales | Virus | Saudi Arabia | 160 |
| (Farzadfar et al., 2006) | Agricultural | Caryophyllales | Virus | Iran | 64 |
| (Freeman et al., 2013) | Agricultural | Fabales | Virus | Australia | 204 |
| (Gauhl et al., 1999) | Agricultural | Zingiberales | Virus | Nigeria | 27 |
| (Gibbs et al., 1999) | Natural | Fagales | Eukaryotic parasite | United Kingdom | 7 |
| (Guadie et al., 2019) | Agricultural | Poales | Virus | Ethiopia | 33 |
| (Guajardo et al., 2019) | Agricultural | Fagales | Eukaryotic parasite | Chile | 5 |
| (Hanna et al., 2008) | Agricultural | Vitales | Virus | Lebanon | 28 |
| (Hatting et al., 2011) | Agricultural | Fabales | Pest | South Africa | 58 |
| (Hay et al., 1992) | Agricultural | Rosales | Virus | New Zealand | 64 |
| (Hillocks et al., 2002) | Agricultural | Malpighiales | Virus | Mozambique | 28 |
| (Hong et al., 2020) | Agricultural | Caryophyllales | Eukaryotic parasite | USA | 29 |
| (Ilbağı et al., 2007) | Agricultural | Poales | Virus | Turkey | 7 |
| (Jones et al., 1996) | Agricultural | Solanales | Virus | United Kingdom | 7 |
| (Karuppaiyan et al., 2015) | Agricultural | Poales | Eukaryotic parasite | India | 16 |
| (Kumar et al., 2012) | Agricultural | Rosales | Virus | India | 5 |
| (Kumari et al., 2006) | Agricultural | Poales | Virus | Yemen | 90 |
| (Kumari et al., 2008) | Agricultural | Fabales | Virus | Eritrea | 56 |
| (Latham and Jones, 2003) | Agricultural | Apiales | Virus | Australia | 29 |
| (Legg and Ogwal, 1998) | Agricultural | Malpighiales | Virus | Uganda | 180 |
| (Li et al., 2018) | Agricultural | Poales | Bacteria | China | 22 |
| (Lindblad and Sigvald, 2004) | Agricultural | Poales | Virus | Sweden | 16 |
| (Mahadevakumar et al., 2019) | Agricultural | Asparagales | Eukaryotic parasite | India | 3 |

| | | | | | |
|---|---|---|---|---|---|
| (Makkouk et al., 2001) | Agricultural | Fabales | Virus | Pakistan | 280 |
| (Makkouk et al., 2003) | Agricultural | Fabales | Virus | Iran | 140 |
| (Mallowa et al., 2006) | Agricultural | Malpighiales | Virus | Kenya | 28 |
| (Mawassi et al., 2018) | Agricultural | Apiales | Bacteria | Israel | 1 |
| (McAllister et al., 2008) | Agricultural | Poales | Virus | USA | 13 |
| (Moodley et al., 2019) | Agricultural | Solanales | Virus | South Africa | 42 |
| (Moreno-Pérez et al., 2014) | Natural | Solanales | Virus | Peru | 6 |
| (Msikita et al., 2005) | Agricultural | Malpighiales | Eukaryotic parasite | Benin | 6 |
| (Muedi et al., 2015) | Agricultural | Fabales | Bacteria | South Africa | 24 |
| (Mueller et al., 2012) | Natural | Apiales | Virus | USA | 8 |
| (Mueller et al., 2012) | Natural | Asterales | Virus | USA | 24 |
| (Mueller et al., 2012) | Natural | Capparales | Virus | USA | 2 |
| (Mueller et al., 2012) | Natural | Caryophyllales | Virus | USA | 14 |
| (Mueller et al., 2012) | Natural | Cucurbitales | Virus | USA | 4 |
| (Mueller et al., 2012) | Natural | Fabales | Virus | USA | 68 |
| (Mueller et al., 2012) | Natural | Gentianales | Virus | USA | 12 |
| (Mueller et al., 2012) | Natural | Lamiales | Virus | USA | 4 |
| (Mueller et al., 2012) | Natural | Malpighiales | Virus | USA | 8 |
| (Mueller et al., 2012) | Natural | Malvales | Virus | USA | 4 |
| (Mueller et al., 2012) | Natural | Poales | Virus | USA | 4 |
| (Mueller et al., 2012) | Natural | Solanales | Virus | USA | 14 |

| | | | | | |
|---|---|---|---|---|---|
| (Mujica et al., 2013) | Agricultural | Vitales | Virus | Chile | 15 |
| (Mulenga et al., 2016) | Agricultural | Malpighiales | Virus | Zambia | 12 |
| (Mulenga et al., 2018) | Agricultural | Malpighiales | Virus | Zambia | 7 |
| (Munck et al., 2018) | Natural | Pinales | Eukaryotic parasite | USA | 4 |
| (Mustafayev et al., 2011) | Agricultural | Fabales | Virus | Azerbaijan | 77 |
| (Naseri and Marefat, 2008) | Agricultural | Fabales | Eukaryotic parasite | Iran | 222 |
| (Ndyetabula et al., 2016) | Agricultural | Malpighiales | Virus | Tanzania | 36 |
| (Nolt et al., 1992) | Agricultural | Malpighiales | Virus | Colombia | 30 |
| (Okao-Okuja et al., 2004) | Agricultural | Malpighiales | Virus | Guinea, Senegal | 10 |
| (Omar and Foissac, 2012) | Agricultural | Cucurbitales | Bacteria | Egypt | 2 |
| (Omar and Foissac, 2012) | Agricultural | Solanales | Bacteria | Egypt | 6 |
| (Oppong et al., 2015) | Agricultural | Poales | Virus | Ghana | 65 |
| (Osti and Marco, 2014) | Agricultural | Ericales | Eukaryotic parasite | Italy | 45 |
| (Ozturk et al., 2008) | Agricultural | Fagales | Virus | Turkey | 11 |
| (Perryman et al., 2009) | Agricultural | Linales | Eukaryotic parasite | United Kingdom | 34 |
| (Phiri et al., 2001) | Agricultural | Gentianales | Eukaryotic parasite | Malawi | 20 |
| (Pongener and Daiho, 2016) | Agricultural | Alismatales | Eukaryotic parasite | India | 20 |
| (Poudel et al., 2019) | Agricultural | Solanales | Virus | USA | 2 |
| (Pourrahim et al., 2007) | Agricultural | Solanales | Virus | Iran | 99 |
| (Prendeville et al., 2012) | Natural | Cucurbitales | Virus | USA | 160 |
| (Rogov et al., 1992) | Agricultural | Caryophyllales | Virus | Kyrgyzstan, Kazakhstan, Uzbekistan | 54 |

| | | | | | |
|---|---|---|---|---|---|
| (Rosso et al., 1994) | Natural | Pinales | Eukaryotic parasite | Argentina | 5 |
| (Sánchez-Campos et al., 1999) | Agricultural | Solanales | Virus | Spain | 24 |
| (Schultz et al., 2015) | Agricultural | Myrtales | Eukaryotic parasite | Brazil | 12 |
| (Sétamou et al., 2000) | Agricultural | Poales | Pest | Benin | 16 |
| (Sétamou et al., 2005) | Agricultural | Poales | Pest | USA | 10 |
| (Shah et al., 2006) | Agricultural | Fabales | Virus | USA | 63 |
| (Silveira et al., 1998) | Agricultural | Solanales | Bacteria | Brazil | 1 |
| (Singh et al., 2019) | Agricultural | Asterales | Eukaryotic parasite | India | 176 |
| (Sisterson et al., 2012) | Agricultural | Rosales | Bacteria | USA | 5 |
| (Soler et al., 2010) | Agricultural | Solanales | Virus | Spain | 21 |
| (Southwell et al., 2003) | Agricultural | Poales | Eukaryotic parasite | Australia | 10 |
| (Springer, 2009) | Natural | Malpighiales | Eukaryotic parasite | USA | 40 |
| (Sseruwagi et al., 2005) | Agricultural | Malpighiales | Virus | Rwanda | 12 |
| (Stanosz et al., 2007) | Agricultural | Pinales | Eukaryotic parasite | USA | 6 |
| (Stevanović et al., 2019) | Agricultural | Rosales | Eukaryotic parasite | Serbia | 23 |
| (Stobbs et al., 2005) | Agricultural | Celastrales | Virus | Canada | 3 |
| (Stobbs et al., 2005) | Agricultural | Juglandales | Virus | Canada | 3 |
| (Stobbs et al., 2005) | Agricultural | Lamiales | Virus | Canada | 1 |
| (Stobbs et al., 2005) | Agricultural | Rosales | Virus | Canada | 31 |
| (Türkölmez et al., 2019) | Agricultural | Solanales | Eukaryotic parasite | Turkey | 7 |
| (Van Vianen et al., 2013) | Natural | Brassicales | Virus | New Zealand | 15 |
| (van Leur and Kumari, 2011) | Agricultural | Fabales | Virus | Australia | 75 |

| | | | | | |
|---|---|---|---|---|---|
| (Were et al., 2013) | Agricultural | Solanales | Virus | Kenya | 30 |
| (Williams et al., 2020) | Natural | Fagales | Eukaryotic parasite | Canada | 39 |
| (Wilson, 1998) | Natural | Asterales | Virus | Australia | 7 |
| (Wilson, 1998) | Natural | Brassicales | Virus | Australia | 5 |
| (Wilson, 1998) | Natural | Caryophyllales | Virus | Australia | 9 |
| (Wilson, 1998) | Natural | Fabales | Virus | Australia | 5 |
| (Wilson, 1998) | Natural | Geraniales | Virus | Australia | 2 |
| (Wilson, 1998) | Natural | Malvales | Virus | Australia | 2 |
| (Wilson, 1998) | Natural | Poales | Virus | Australia | 3 |
| (Wilson, 1998) | Natural | Ranunculales | Virus | Australia | 1 |
| (Wilson, 1998) | Natural | Solanales | Virus | Australia | 1 |
| (Wraight et al., 1993) | Agricultural | Poales | Pest | USA | 12 |
| (Wu and Subbarao, 2006) | Agricultural | Asterales | Eukaryotic parasite | USA | 302 |
| (Wu and Subbarao, 2006) | Agricultural | Capparales | Eukaryotic parasite | USA | 4 |
| (Wu and Subbarao, 2006) | Agricultural | Fabales | Eukaryotic parasite | USA | 1 |
| (Wu and Subbarao, 2006) | Agricultural | Solanales | Eukaryotic parasite | USA | 1 |
| (Wydra and Verdier, 2002) | Agricultural | Malpighiales | Bacteria | Ghana, Benin | 8 |
| (Wydra and Verdier, 2002) | Agricultural | Malpighiales | Virus | Ghana, Benin | 8 |
| (Yahaya et al., 2019) | Agricultural | Poales | Virus | Nigeria | 2 |
| (Yahaya et al., 2019) | Natural | Poales | Virus | Nigeria | 1 |
| (Yaninek et al., 1996) | Agricultural | Malpighiales | Pest | Benin | 6 |

*Climate and weather data*

After extracting relevant data from each study, we paired each plant disease observation with climate reanalysis data extracted using Google Earth Engine (Gorelick et al., 2017). First, we specified circular buffers around the centroid of each observation with diameter equal to the approximate spatial distance of the survey.

We then extracted temperature and precipitation (weather) data from the ERA5-land monthly averaged ECMWF climate reanalysis dataset (Sabater, 2019) for the location and month(s) in which each incidence survey occurred, and calculated the mean temperature and mean daily precipitation over the months of the survey period. The ERA5-land monthly dataset uses a combination of models and observations to provide climate and weather data for areas on land, spanning 1981–2022. The spatial resolution of the dataset is 11.1km. Next, we extracted historical average temperature and precipitation data for the location of each observation from the WorldClim BIO Variables database (Hijmans et al., 2005). This dataset represents the 30-year average temperature and precipitation in a location from between 1960-1990, with a spatial resolution of 0.9km.

### Additional details on analyses

*Data processing*

Our systematic literature search led to 5906 observations from 129 unique studies. We then removed 193 of these observations because we subsequently determined that either the survey occurred in water, the scale of the survey was not at the at the level of the individual plant (e.g., they were surveying different pinecones on the same tree for disease), or because the survey did not fall into either our "wild" or "agricultural" categories. We then removed 333 observations because they lacked climate or weather data. Our final database was therefore 5380 observations from 118 unique studies. Prior to analyses, we standardized historical and contemporaneous temperature and precipitation data as z-scores in R (R Core Team, 2022).

*Correlated data*

In our database, contemporaneous temperature and historical temperature were positively correlated ($r = 0.564$; Fig. S1), as were precipitation and historical precipitation ($r = 0.560$; Fig. S1). Correlations between either temperature variable with either precipitation variable were weak (Fig. S1).

The positive correlations between contemporaneous and historical temperature and precipitation can be observed in the underlying data (Fig. S2). For example, our database did not include any surveys of natural plant populations that were from a historically warm climate (e.g. >20°C) that also had a relatively cool contemporaneous temperature (e.g. <20°C) at the time of the survey (Fig. S2a). As discussed in the main text, these data features can make it more difficult to extrapolate predicted effects, for example in natural systems from cold temperatures to warm temperatures, or from warm to cold temperatures in agricultural systems. Experimental temperature work can supplement observational approaches by overcoming these constraints, testing both natural and agricultural plant systems from different historical climates against a range of experimental temperatures.

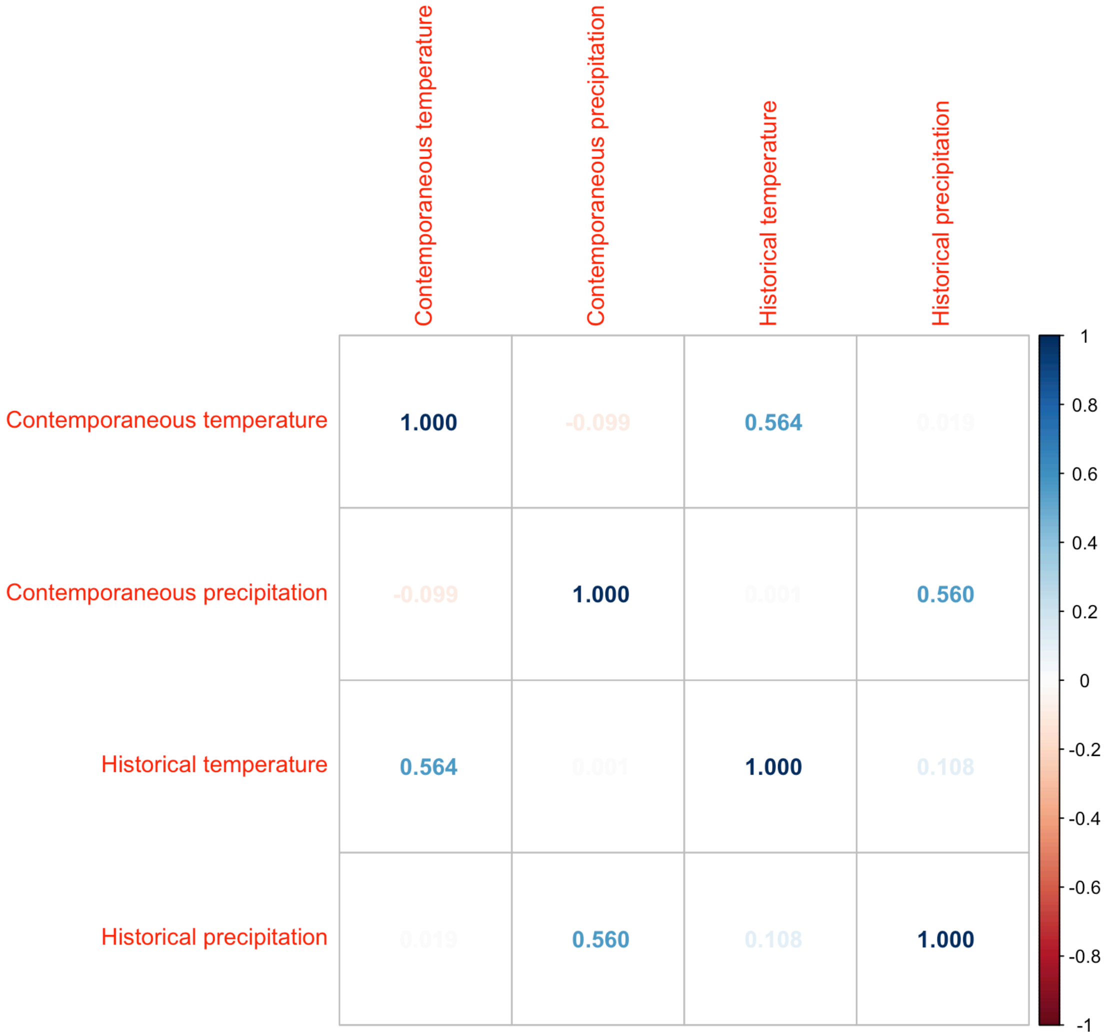


Fig. S1. Correlation matrix for the two weather (contemporaneous temperature and precipitation) and two climate (historical temperature and precipitation) variables. Darker blue values represent larger positive correlations, while faint values represent weaker correlations closer to 0.

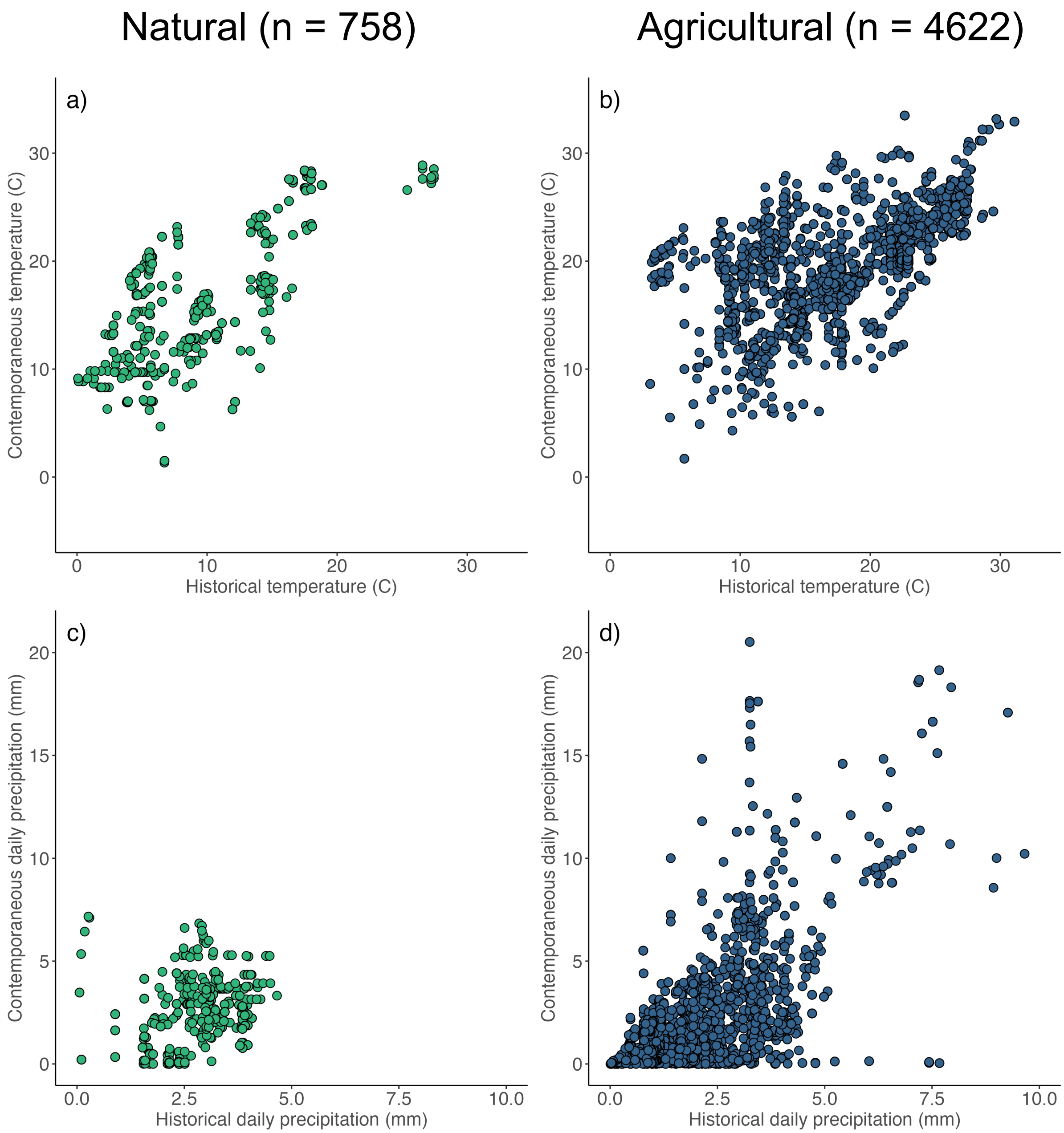


Fig. S2. Scatterplot of underlying temperature (a,b) and precipitation (c,d) data in natural (a,c) and agricultural (b,d) systems.

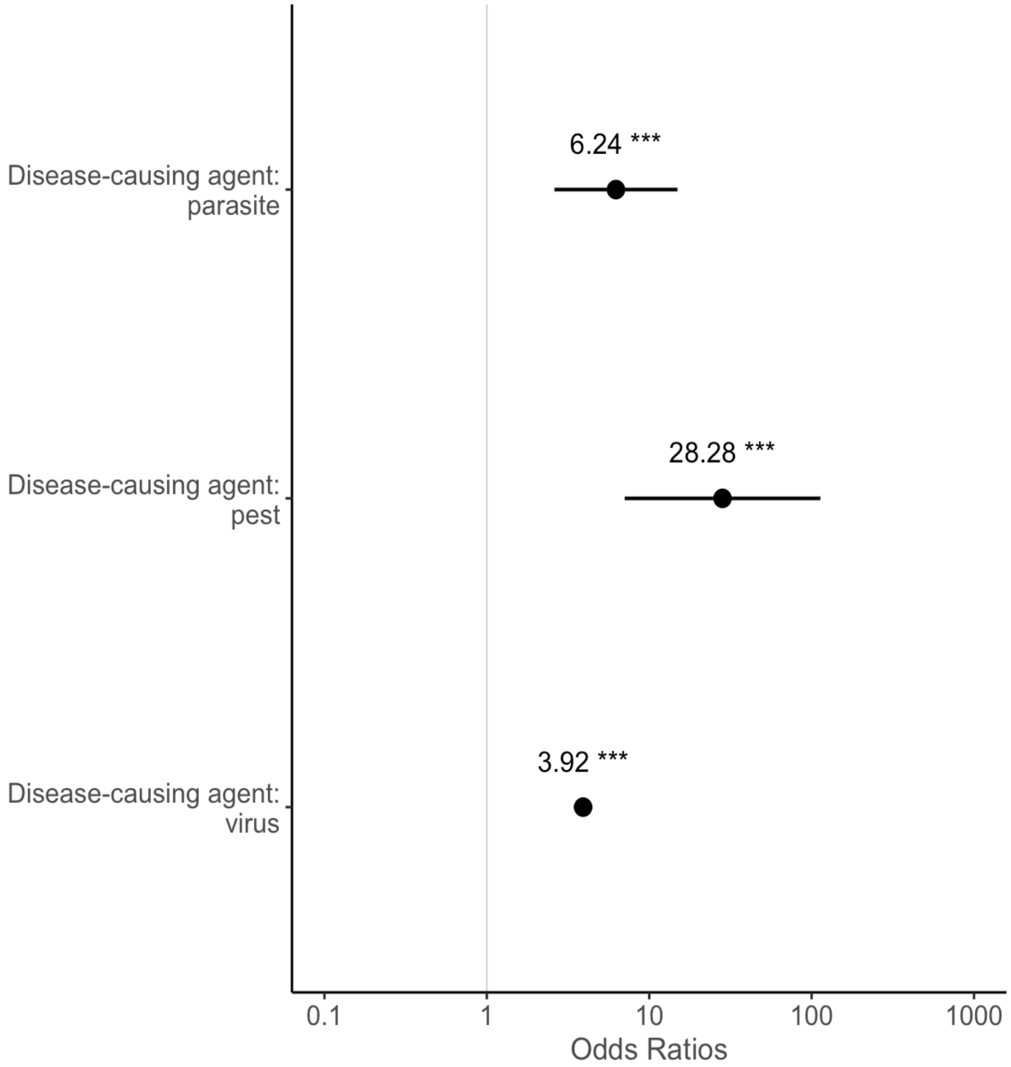


**Fig. S3. Plant disease risk varied by type of disease-causing agent.** Points and error bars represent estimated odds ratios and 95% confidence intervals, respectively, where higher odds ratios represent greater disease risk. Odds ratios are expressed relative to the fourth type of disease-causing agent included in our model: bacteria, which was associated with the lowest risk of disease incidence. Odds ratios associated with climate and weather factors included in the model are shown in Fig. 2 of the main text.

*Secondary analyses*

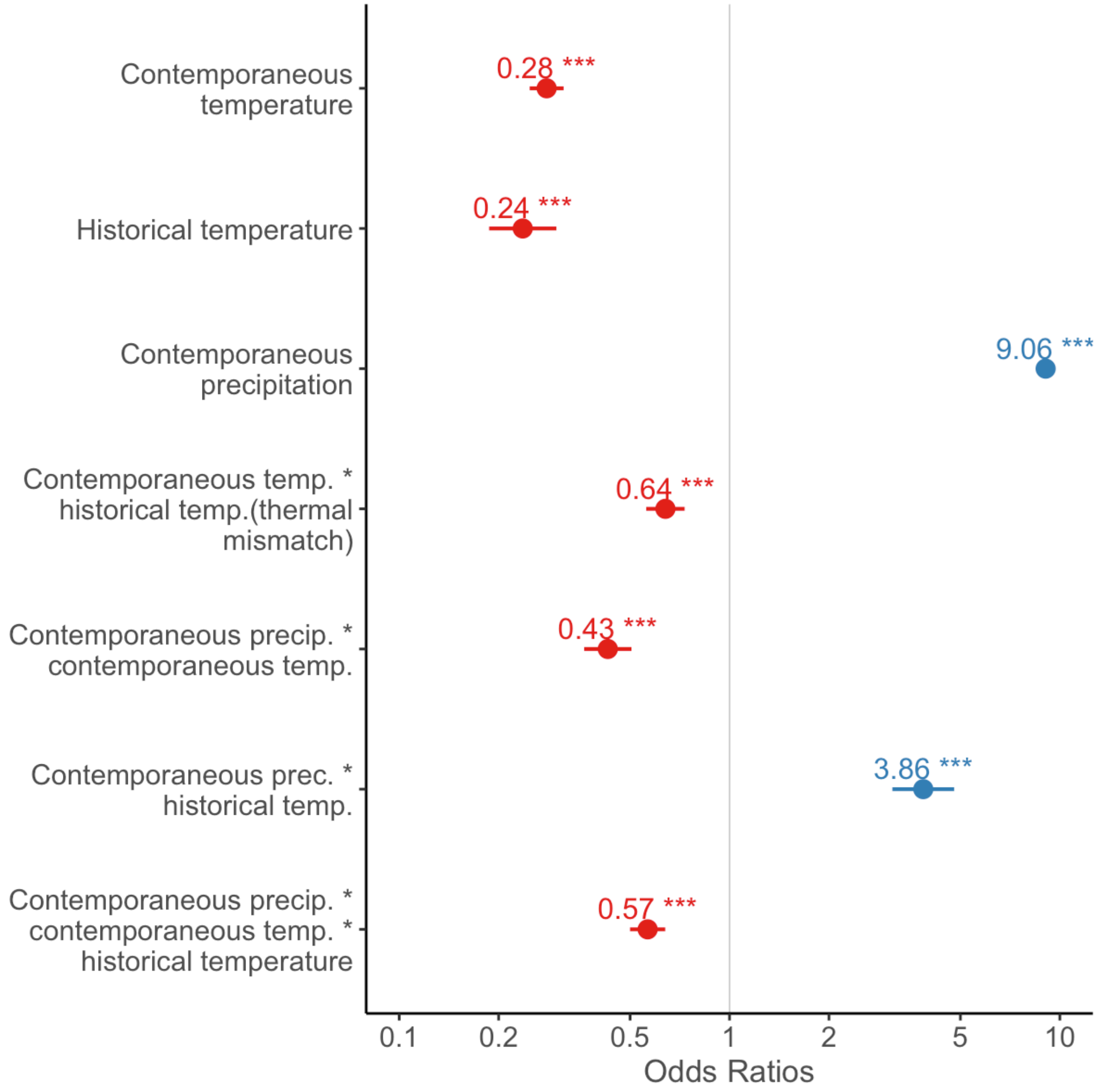


**Fig. S4. Thermal mismatches become stronger (more negative) in wild systems with high contemporaneous precipitation.** Effect sizes of modeled variables from a secondary analysis for wild systems. Points and error bars represent estimated odds ratios and 95% confidence intervals, respectively, where higher odds ratios represent greater disease risk. Blue points show odds ratios greater than 1, with red points showing odds ratios less than 1. Contemporaneous and historical temperature and contemporaneous precipitation were standardized before model fitting. Thermal mismatches represent an interaction between contemporaneous temperature by historical temperature, and the three-way contemporaneous precipitation by contemporaneous temperature by historical temperature interaction represents the effect of precipitation on thermal mismatches—the key finding of this analysis.

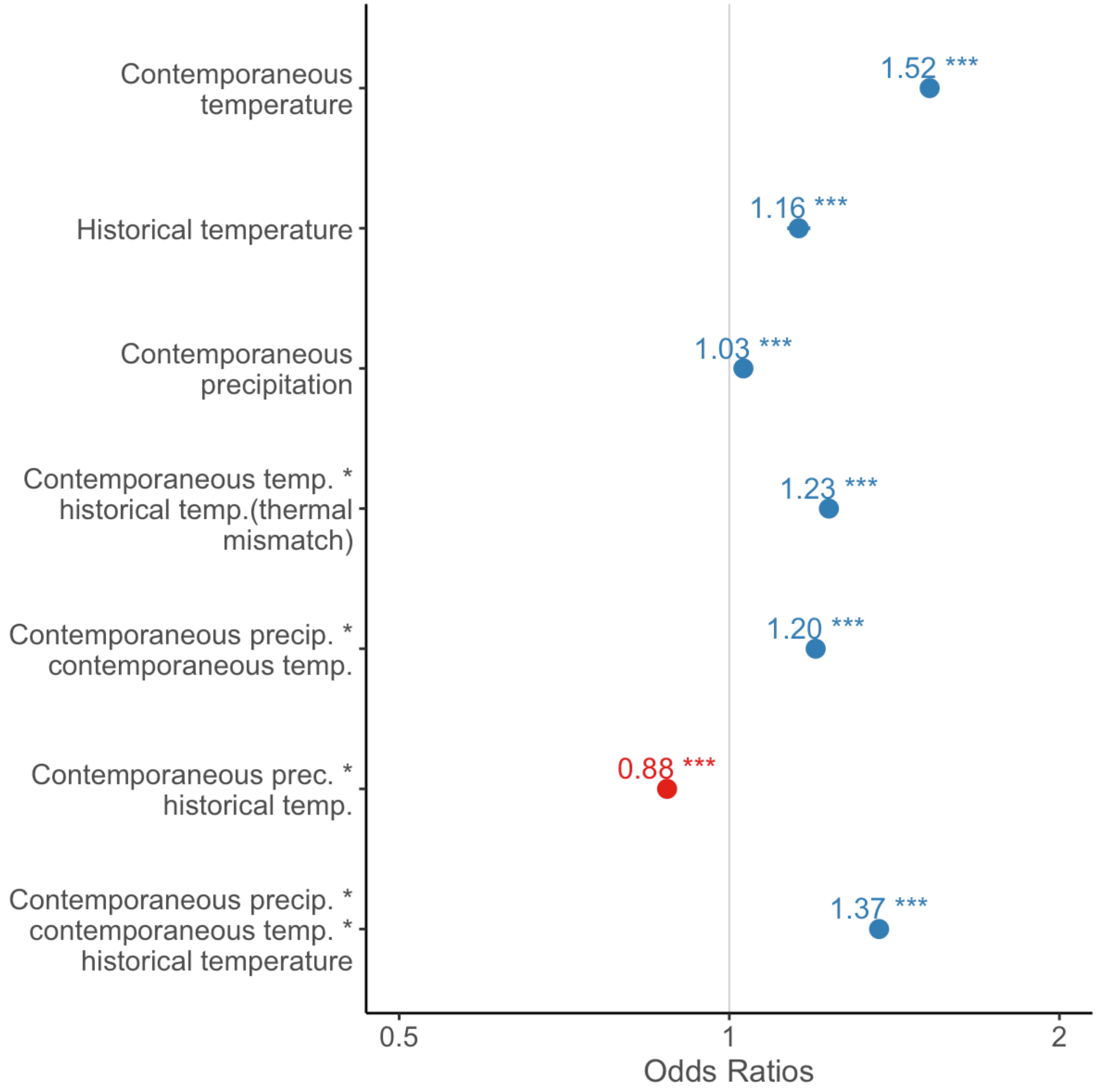


**Fig. S5. Effects of warm weather are exacerbated in agricultural systems with wet weather and warm historical temperatures.** Effect sizes of modeled variables from a secondary analysis for agricultural systems. Points and error bars represent estimated odds ratios and 95% confidence intervals, respectively, where higher odds ratios represent greater disease risk. Blue points show odds ratios greater than 1, with red points showing odds ratios less than 1. Contemporaneous and historical temperature and contemporaneous precipitation were standardized before model fitting. Thermal mismatches represent an interaction between contemporaneous temperature by historical temperature, and the three-way contemporaneous precipitation by contemporaneous temperature by historical temperature interaction represents the effect of precipitation on thermal mismatches.
